# Transcranial alternating current stimulation (tACS) mechanisms and protocols

**DOI:** 10.1101/138834

**Authors:** Amir Vala Tavakoli, Kyongsik Yun

**Affiliations:** Computation and Neural Systems, California Institute of Technology, Pasadena, CA 91125, USA; Division of Biology and Biological Engineering, California Institute of Technology, Pasadena, CA 91125, USA; Department of Psychology, University of California, Los Angeles, CA, USA; Bio-Inspired Technologies and Systems, Jet Propulsion Laboratory, California Institute of Technology, Pasadena, CA 91125

**Keywords:** Neuromodulation, entrainment, oscillatory, review, tACS, EEG, tES, NIBS, NTBS

## Abstract

What if you could affect both neuroplasticity and human cognitive performance by parametrically modulating neural oscillations? Ongoing neuronal activity is susceptible to the modulation of synaptic activity and membrane potentials. This susceptibility leverages transcranial alternating current stimulation (tACS) for neuroplastic interventions. Through neuromodulation of phasic, neural activity, tACS presents a powerful tool for investigations of the neural correlates of cognition alongside other forms of transcranial electric stimulation (tES) and noninvasive brain stimulation (NIBS). The rapid pace of development in this area requires clarification of best practices. Here, we briefly introduce tACS dogma and review the most compelling findings from the tACS literature to provide a starting point for the use of tACS under research conditions.

## 1. Introduction

Technological and ethical constraints have forced investigations of human cognition to rely upon usually non-invasive electrophysiology and neuroimaging techniques to reveal the neural correlates of perceptual, cognitive and behavioral functions. Investigating the causal neurophysiology subtending function requires recent methodological innovations to advance neuroscience. Transcranial alternating current stimulation (tACS) has emerged as a unique tool, given its particular advantages. An evolution on clinical transcranial electric current stimulation (tES), tACS is a non-invasive technology that helps engender neuroscientific results with greater inferential force.

Consensus on tACS methods have been slow to emerge. While the absence of a methodological gold-standard risks unreliable results and hinders systematic improvement of the field, rigorous tACS methods have emerged. This review’s criterion for inclusion of scholarship considered a growing agreement on the importance of controls: principally, subject-blinding and either sham or active sham conditions. More detailed reviews of tACS methods have been compiled elsewhere (Neuling et al., 2016; Schutter & Wischnewski, 2016; Veniero et al., 2015). This mini-review is intended to highlight: 1. key advantages; 2. assumptions; and 3. protocols.

## 2. The advantages of tACS: The tACS current, physiological control, and causality

### 2.1 The recent expansion of neuromodulation research

Recent interest in neuromodulation traces partly to the desire to investigate cognitive function in a parametrically rigorous manner. Unlike other forms of neuromodulation, tACS enables the manipulation and entrainment of intrinsic oscillations through sinusoidal currents (Antal & Paulus, 2013; Paulus, 2011; Thut et al., 2011). The phasic profile of a tACS current alternates regularly between a positive and negative voltage. Alternatively, in transcranial direct current stimulation (tDCS), the current describes a monophasic, distinctly non-oscillating baseline voltage. Endogenous activity is either modulated by depolarization (anodal) or hyperpolarization (cathodal), in a global flow of current that supplies electrons to the anodal electrode (promoting endogenous oscillations) and retracts electrons from the cathodal electrode (suppressing endogenous oscillations) (Song et al., 2014).

In tACS, an oscillating current rhythmically reverses electron flow. Averaged over an extended temporal-window, unlike tDCS, the tACS current omits any directional voltage component. In oscillating tDCS (otDCS), oscillations ride a directional component (Guleyupoglu et al., 2014). Transcranial random noise stimulation (tRNS) is used to inject an alternating current of bounded stochasticity (Saiote et al., 2013). While different stimulation protocols have not fully clarified the neurophysiological mechanism of each method, it is now settled that oscillatory states anticipate cognitive phenomena (Schutter & Wischnewski, 2016; Donner & Siegel, 2011; Wang, 2010).

### 2.2 Physiological advantages (parametric modulation of neural oscillations)

The most prominent mode of neuroplasticity, long-term potentiation (LTP) (Lee and Silva, 2009), is informed by spike-time dependent neurophysiology. One fundamental advantage of tACS is the physiological entrainment of large populations of cells. During entrainment, an endogenous oscillation synchronizes with exogenous, rhythmic stimulation. Investigators have effectively used rhythmic photic stimulation to assay visual cortices for susceptibility to entrainment (Adcock et al., 2012; Poleon & Szaflarski, 2017; Adrian & Matthews, 1937). This suggests that frequency and phase information are fundamental variables in neural function (Herrmann et al., 2016). tACS can bypass sensory stimulation entirely, by instead inducing entrainment through an exogenously applied, nearly imperceptible alternating current. Clarifying the causal relationship between cognitive function and oscillatory activity will require the combination of behavioral (Kanai et al. 2008; Feurra et al. 2011; Laczó et al. 2012) and electrophysiological (Zaehle et al. 2010) methods (Thut, 2011; Herrmann et al., 2014). This joint approach requires the testing of clear physiological assumptions.

tACS is capable of parametrically modulating neurophysiology. Endogenous oscillations are constituted by interactions with oscillatory inputs from near and eccentric neural sources (Herrmann et al., 2016; Romei et al., 2016; Vosskuhl et al., 2016). These aggregate neural oscillations dynamically stabilize to form a set of functional bands of activity that are recordable with EEG and susceptible to entrainment (Herrmann et al., 2016; Romei et al., 2016; Thut et al., 2011). It remains an open question whether our neurology is predisposed to replicate natural frequencies in the environment, or if exogenous frequencies alter connectivity through classical mechanisms of spike time dependent plasticity (STDP), or both (Kasten et al., 2016; Vossen et al., 2015; Zaehle et al., 2010).

**FIGURE 1.**
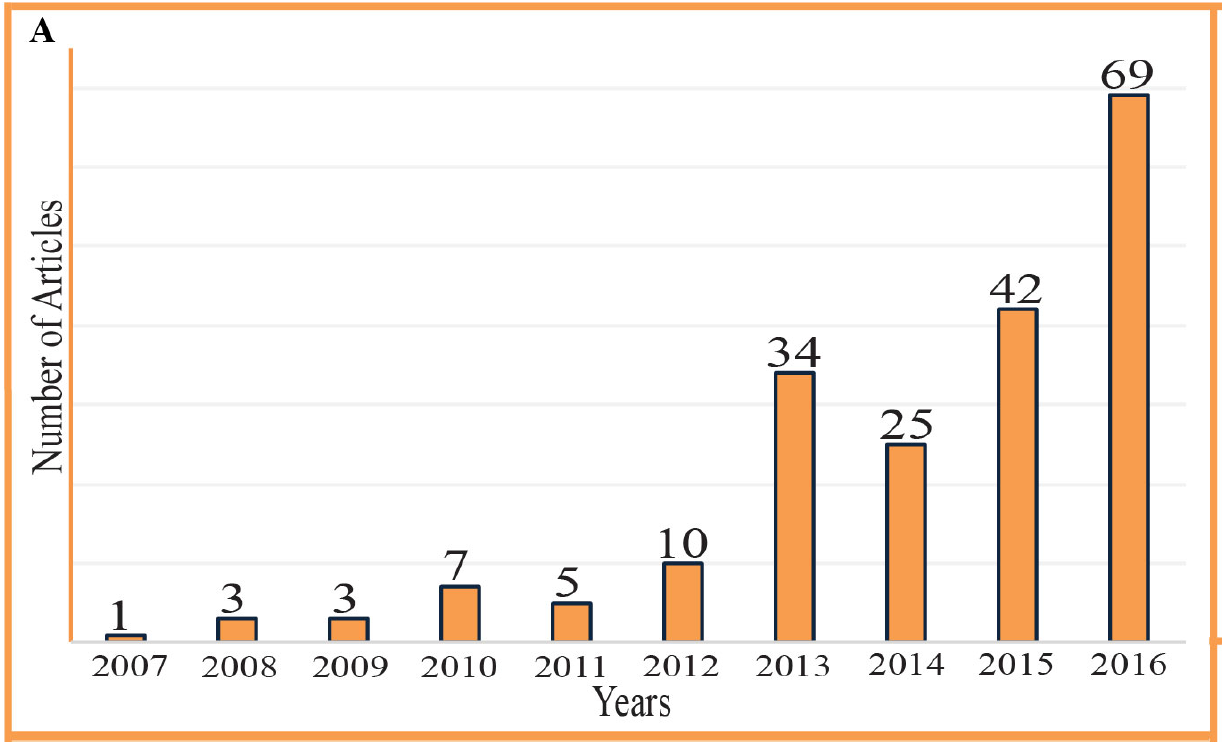
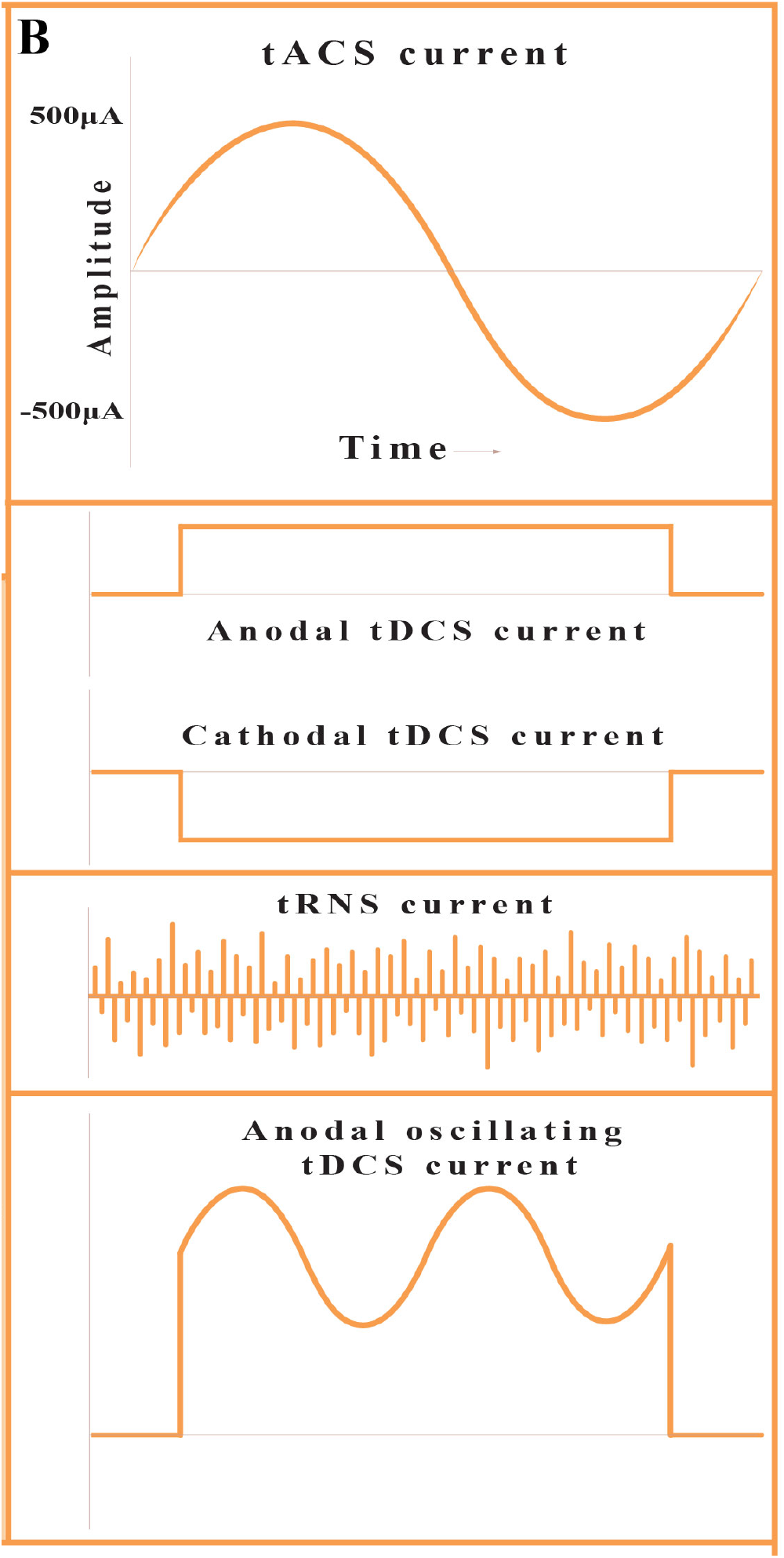
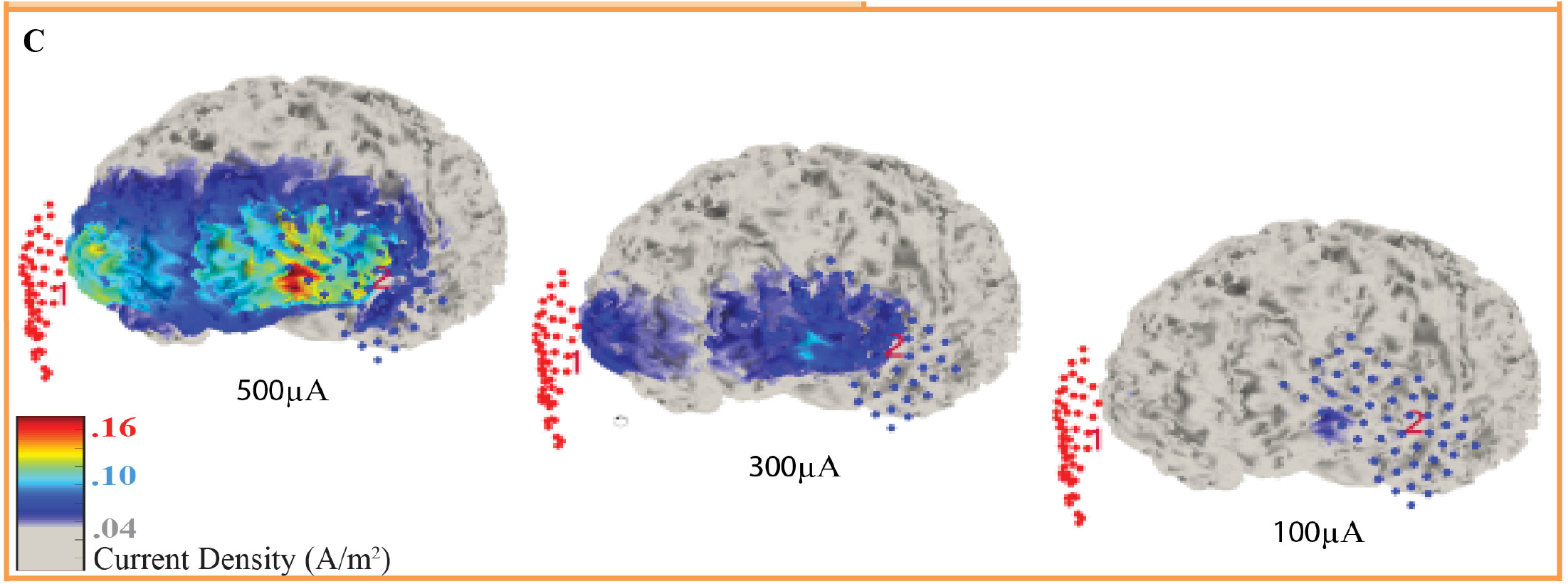
Recent growth and the tACS current profile. **(A)** Number of transcranial alternating current stimulation studies in the last 10 years. PubMed listed articles that used the term “transcranial alternating current stimulation” in the title or in the abstract were counted. The technique has been increasingly applied in the recent years and we may predict the exponential increase in the number of studies in the upcoming years. **(B)** Transcranial current stimulation protocols. *tACS (transcranial alternating current stimulation), tDCS (transcranial direct current stimulation), tRNS (transcranial random noise stimulation), otDCS (oscillating transcranial direct current stimulation). **(C)** Computational modeling of cortical current density while stimulating with tACS. Scm X Scm electrodes were placed on the F3 and F4. Three brains represent 500μA, 300μA, and 100μA stimulations. The size of the affected cortical regions and the current density both increased as the stimulation increased. Stimulation intensity and electrode size should be carefully determined based on the size of the target region (Lee et al., 2017).

### 2.3 tACS facilitates potentially causal inferences

Frequently, (dependent) electrophysiological variables have indexed (independent) cognitive processes (Herrmann et al., 2016). NIBS methods enable investigators to reverse these dependencies where observational-correlational methods cannot (Poldrack, 2006). Furthermore, EEG research has covered much terrain in the oscillatory phenomenology of cognition and behavior. Well defined cognitive functions are commonly attributed to a subset of specific oscillatory features and frequencies. Buoyed by such a priori electrophysiological evidence, investigators use tACS to extend causal explanations of (independent) electrophysiological variables to (dependent) cognitive processes. Thus, measured behavior becomes a function of parameterized electrophysiology.

### 2.4 Recent focus on determining behavioral causality from neurophysiology

Causal interpretations of neural systems and functional circuitry demand neuromodulation at multiple scales of cortical network activity (Ruffini et al., 2014). While the imaging of voltage-related neurophysiology with multi-photon microscopy is still nascent, optogenetics and invasive electrophysiology (Fröhlich & Schmidt, 2013; Kuki et al., 2013; Anastassiou & Koch, 2015) enable causal investigations of large-scale dynamics in animal models. Invasive investigation through human electrophysiology, however, have determined local entrainment to alternating currents (Amengual et al., 2017). Clinical pursuit of effective connectivity has targeted functional circuits by registering intracranial oscillations to tractography (Elias et al., 2012). Functionally, deep brain stimulation of the human hippocampus has revealed the effect of 50 Hz currents on memory performance (Ezzyat et al., 2017). Noninvasively, neuroplasticity is being induced across multiple functional domains (Hameed et al., 2017; Bolognini et al., 2009). Also, tACS appears more effective than tDCS for network entrainment (Ali et al., 2013) and, in an animal model of epilepsy, closed-loop tACS appears to mitigate spike-wave effects (Berényi et al., 2012). Thus, investigations of neural systems are active across a range of scales, models and contexts.

### 2.5 EEG-tACS and feedback-controlled studies enable stronger inferences

Feedback-control enables precise control of exogenous and endogenous oscillations. Thus tACS is now combined with other NIBS, as well as EEG, to sharpen experimental inferences (Boyle & Fröhlich, 2013; Lustenberger et al., 2016; Raco et al., 2016; Roh et al., 2014; Kanai et al., 2010). While many EEG-tACS experiments have recorded subjects’ pre- and post-stimulation EEG to avoid signal artifacts (Veniero et al., 2015; Zaehle et al., 2010), recent refinements to simultaneous EEG-tACS better elucidate endogenous oscillations (Neuling et al., 2017; Roh et al., 2014). While EEG-tACS enables sub-millisecond temporal resolution (tenOever S., 2016; Neuling et al., 2015), concerns about signal artifacts persist (Noury et al., 2016). Closed-loop tACS-TMS protocols also enable parametric adjustment of magnetic stimulation to instantaneous physiology (Thut et al., 2017). Raco et al. (2016) developed a closed-loop protocol wherein instantaneous tACS-phase triggered TMS pulses. While some research questions do not demand online protocols, future combination of these techniques may enable stronger causal attributions to oscillations (Frohlich; Herrmann et al., 2016).

**Figure 2.**
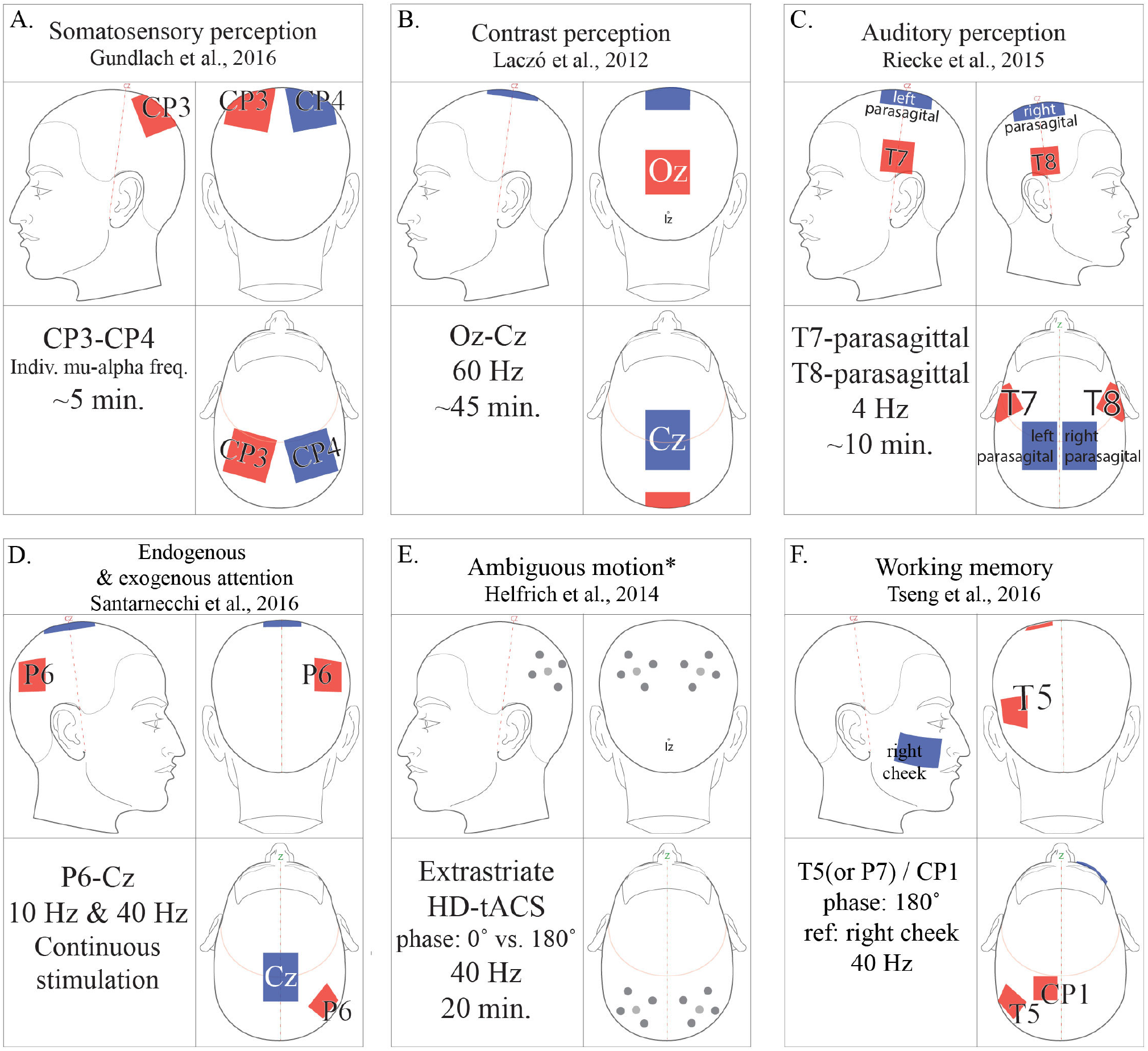
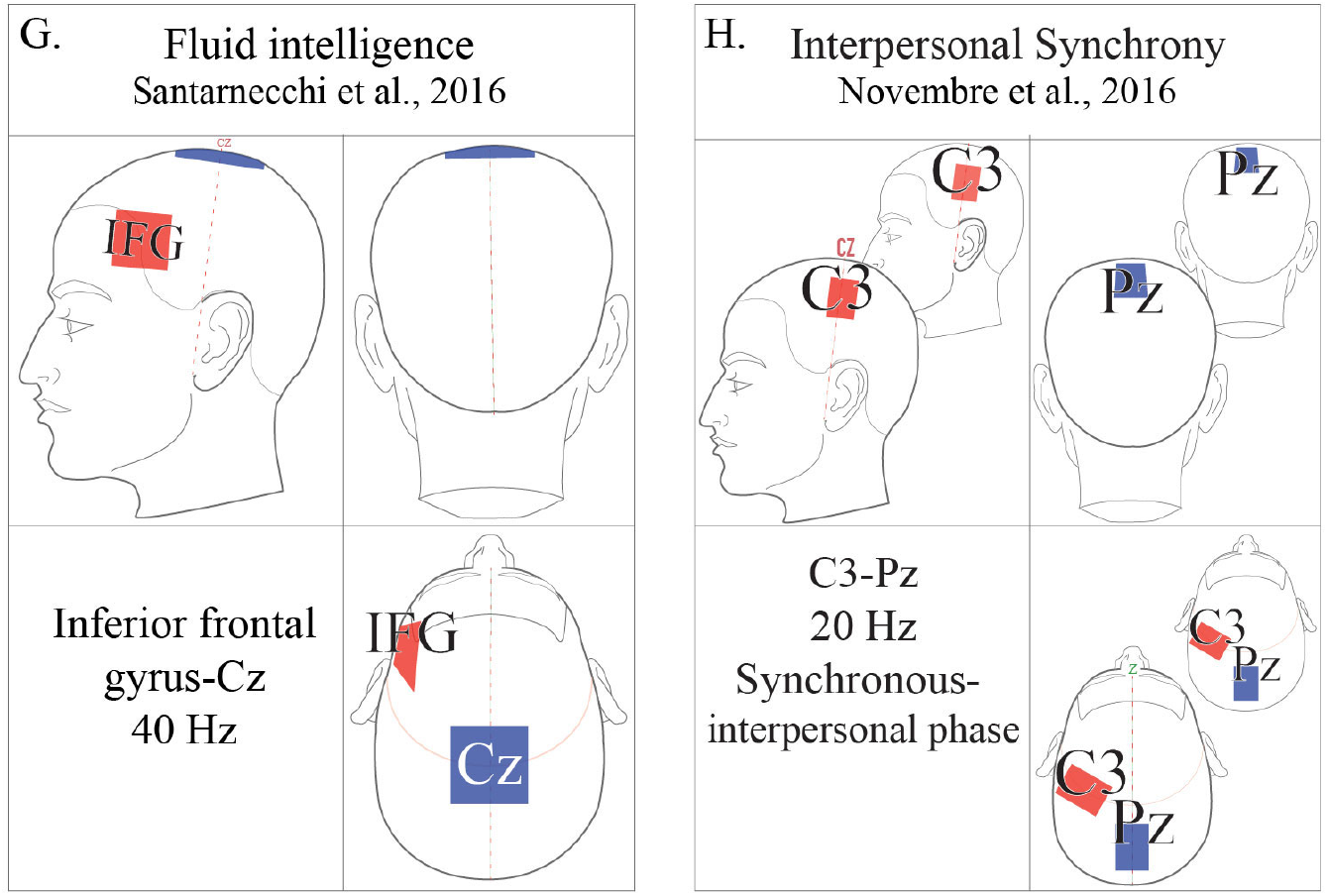
tACS protocols. (A) 1000 μA; (B) 1500μA; (C) >1000 μA; (D) 2000 μA; (E) Amplitude thresholded; (F) 1500 μA; (G) 750 μA; (H) 1000 μA. All amplitudes in peak-to-peak microamps. *Ambiguous motion protocol alternates bothphase-offset and HD-montage.

## 3. Methodological considerations for rigorous tACS experiments

The methods described in this review represent early forays into neuromodulation using tACS. Valid results require that all modes of NIBS be performed while taking precautions to increase replicability and validity.

Subject-awareness of condition assignment must be avoided through established blinding methods. While tACS emits no sound to cue stimulation, blinding should be enforced by subtending subject-specific stimulus detection thresholds for both visual and somatic percepts. Rostral electrode montages induce phosphenes more easily and at lower amplitudes (Schutter, 2016; Neuling, 2012?). With active shamming, broader comparisons are enabled by changing montage or frequency while constraining all remaining parameters (Fuerra et al., 2011; Mehta et al., 2015; Schutter, 2016). Both sham and experimental currents should integrate comparable ramping periods (Woods et al., 2016). Future investigations may additionally facilitate replicability by implementing double-blinding through fully automated tACS protocols.

Given the neuro-oscillatory effects of visual information, context is particularly important in tACS experiments, where subjects’ endogenous states are the focus of investigation (Reato et al., 2013). Lighting conditions can influence detection-thresholds for tACS-induced phosphenes (Kanai et al., 2008; Paulus, 2010; Neuling, 2013). While some suggest that tACS electric fields fall below published thresholds of retinal sensitivity, challengers suggest that the dark adaptation of the retina contributes to the frequency at which retinal excitation occurs (Herrmann, 2013; Kanai et al., 2008; Neuling, 2013; Paulus, 2010;). tACS can also modulate individual alpha frequency (IAF). Eyes-closed EEG states are marked by high baseline alpha-band power (Herrmann et al., 2016). Thus, predictably, Neuling et al. (2013) used within-band tACS to increase IAF power, specifically for eyes-open conditions. Replicable tACS results thus require investigators to treat neuromodulation as a function of contextual predictors of ongoing brain states.

While tACS protocols vary significantly, general refinements, optimal protocols and function-specific parameters have emerged (Frohlich). To avoid visual artifacts while targeting rostral cortical regions, investigators can improve the locality of currents with “ring” electrode montages. Here, a single stimulation electrode is encircled by four reference electrodes (Helfrich et al., 2014). Many experiments predetermine stimulation frequency for all subjects (Moisa et al., 2016; Riecke, 2016). Between-subject variability in the bounds of EEG bands, such as 7-12 Hz alpha, have precipitated subject-specific stimulation frequencies, determined by peak band power (Mehta et al., 2015; Herrmann et al., 2016). Reference electrode montage is also crucial to achieve desired current densities and stimulation. Mehta et al. (2015) compared entrainment of peak physiological tremor with contralateral reference electrodes as well as extracephalic electrodes placed on either ipsilateral or contralateral shoulders. They determined that only the contralateral extracephalic reference montage entrained peak physiological tremor (Mehta et al., 2015). Additionally, multi-electrode montages allow multiple currents to be applied in- or out-of-phase, enabling investigation of inter-hemispheric coherence (Struber et al., 2014).

Another methodological dimension is subject experience. Before initiating an experiment, experimenters should address subjects’ anxieties about the electrical current through brief exposure. Humane precautions may preempt artifacts in both oscillatory and behavioral data, arising from experimentally non-salient autonomic arousal (Bonnet & Arrand, 2001).

Finally, subjects’ safety and the continued refinement of tACS methods require continued vigilance for potential behavioral changes that may persist beyond post-stimulation measurements. Parameter-dependent investigations into these aftereffects continue to emerge (Matsumoto & Ugawa, 2017; Herrmann, 2016; Veniero et al., 2015; Wach et al., 2013; Neuling, 2013; Antal et al., 2008).

## 4. Target and task-specific tACS

### 4.1 Attention

tACS has been of interest to direct investigations of endogenous and exogenous attention. Hopfinger et al., (2016) investigated the effects of alpha and gamma tACS on endogenous and exogenous attention by comparing subjects’ performance on two spatial cueing tasks. In their study, 40 Hz gamma tACS had a facilitative effect on endogenous orienting, but no significant effect on exogenous orienting, suggesting a critical role of low gamma in attentional disengagement and reorientation (Hopfinger et al., 2016).

### 4.2 Perception research

Using EEG, investigators have pursued the oscillatory correlates of perceptual phenomena such as the ventriloquism effect (Kumagai & Mizuhara, 2016), the double-flash illusion (Cecere et al., 2015) and mirrored social embodiment (Oberman et al., 2005; Raymaekers, 2009). Leveraged by common electrophysiology, tACS has demonstrated its utility for perceptual investigations.

There is growing evidence of the consequences of tACS on audition (Riecke, 2016; Baltus & Herrmann, 2016). By applying a 1000 µA DC current, described by an approximately 425 µA, 10 Hz component, Neuling et al., (2012) found a causal relationship between oscillatory phase and auditory signal-detection. While this study was not a pure instantiation of tACS, other investigations of oscillations and audition have demonstrated a functional role of alpha (Weisz et al., 2011) and delta/theta frequencies (Riecke et al., 2015). Speech perception has also been effected with 40 Hz tACS (Rufener et al., 2016_1_; Rufener et al., 2016_2_)

Protocols specifically targeting the visual cortex have faced significant challenges from reviewers despite innovations in tACS montage. While some have suggested posterior montages, for example active at Oz and referenced to Vertex (Cz), purely non-retinal stimulation of the visual cortex remains somewhat contentious (Paulus, 2013; Schutter, 2016). Yet, while the targets and corresponding montages of multiple, effective tACS experiments have applied montages anterior to the occipital cortex, the induction of retinal phosphenes has not emerged as a significant confound of experimental results (Pogosyan et al., 2009; Marshall et al., 2006; Kirov et al., 2009; Kanai et al., 2010). Rigorously considered implementation of tACS currents is compatible with controlled neuroscientific investigations.

In the visual domain, investigators have successfully modulated motion perception (Helfrich et al., 2016; Helfrich et al., 2014; Strüber et al., 2013), mental rotation (Kasten & Herrmann, 2017), visuo-motor coordination (Santarnecchi et al., 2017), and induced phosphenes (Kanai et al., 2008) -- the perception of light that is purely neural and non-photic in origin. The tACS literature pays significant attention to controlling for phosphenes and ensuring that subjects experience experimentally useful phosphenes in a parameterized manner (Schutter, 2016). Care should be taken to determine subject-specific parameters for phosphene induction thresholds and to perform experiments at or below these thresholds (Kanai et al., 2008). Many investigators additionally query subjects for phosphene percepts (Schutter, 2016; Strüber et al., 2014; Antal et al., 2008). Investigators of tACS-induced phosphenes have compared current profiles in light and dark conditions (Schutter, 2016). Kanai et al. (2008) induced qualitative changes in phosphene perception, such as differences in position, orientation, diffusivity, and temporal stability (flickering). In lighted conditions, stimulation in the beta range (20 Hz) induced low phosphene-detection thresholds and qualitatively stronger phosphenes, whereas in dark conditions, stimulation in the alpha range (10 Hz-12 Hz) induced the strongest phosphenes. Thus, the frequency range between 10 to 40 Hz (Paulus et al., 2011; Moliadze et al., 2010a) risks phosphene interference. Beyond phosphenes, Herrmann et al. (2014) demonstrated that the perceived direction of apparent motion can be modulated in an ambiguous motion task by applying bilateral, anti-phase tACS in the gamma band.

### 4.3 Motor function

tACS has been used to investigate motor enhancement, learning and memory. While appropriate montages at C3/C4 in the international 10-20 system target the contralateral limb, motoric regions of interest exist outside of the precentral gyrus. A tACS-fMRI investigation revealed that behavioral changes positively correlated with BOLD activity in M1 but negatively correlated with activity in dorsomedial prefrontal cortex, a region regarded as a locus of executive motor control (Moisa et al., 2016). Brinkman et al. (2016) compared alpha and beta tACS to investigate movement selections. Enhanced movement acceleration and velocity have been achieved with gamma band entrainment over primary motor cortex (M1) (Moisa et al., 2016) and sensorimotor integration with beta tACS (Guerra et al., (2016). It was determined that alpha-band tACS could cause an almost 40 min decrease in corticomuscular coherence, an established measure of functional coupling between M1 and musculature (Wach et al., 2013). While motor learning improved with application of 10 Hz tACS (Antal et al., 2008), with similar results obtained at higher amplitudes (Nitsche, Liebetanz, et al., 2003), motor memory was enhanced by applying tACS during sleep (Fröhlich et al., 2016).

### 4.4 Memory, learning and higher cognition

Investigations of memory, learning, and higher cognitive functions have been attempted. Notably, investigations of forebrain function should consider visual artifacts while using rostral montages. Performance on visual memory-matching tasks compared in-phase, bilateral, theta-band stimulation to anti-phase stimulation. In-phase theta was found to reduce reaction times on a visual memory-matching task, whereas anti-phase degrades performance and increased RTs (Polania et al., 2012). Alekseichuk et al., (2016) found that spatial working memory relies upon theta-gamma, cross-frequency coupling. Also, in an experiment applying feedback-controlled 12 Hz tACS stimulation during sleep, there was no significant increase in declarative memory consolidation, despite increased motor memory consolidation and enhanced sleep spindle activity (Fröhlich et al., 2016). In an investigation of reversal-learning, a task during which subjects are trained on a discrimination task followed by periods of target-distractor reversals, participants receiving theta (6 Hz) stimulation over the frontal cortex experienced faster reversal learning (Wischnewski et al., 2016). They additionally experienced an increase in risk-taking behavior compared to sham participants.

### 4.5 Risk-taking

Interhemispheric phase-difference appears to affect executive decision-making. An investigation of risk-taking using the Balloon Analog Risk Task found that theta (6.5 Hz) stimulation over the left hemisphere is capable of increasing risk-taking behavior (Sela et al., 2012). A bi-frontal, anti-phase protocol used in an investigation of reversal-learning by Wischnewski et al., (2016) should also be noted for an unexpected effect on risk-taking.

## 5 Summary and future directions

Our goal has been to clarify the benefits of tACS to neural investigations. Despite many methodological advances, unraveling the neurophysiological and oscillatory complexity of cognitive function demands investigation at multiple scales.

Investigators studying consciousness through psychophysics suggest the vital role played by systems-level information integration, such as audio-visual integration (Bhattacharya et al., 2002; Shams et al., 2000). While deep-brain integrators of neuronal information have been revealed in animal models (Fetsch et al., 2013), causal determinations have been significantly more difficult in humans (Beauchamp et al., 2004). Yet, the electrophysiological literature suggests the emergence of perception, cognition and consciousness from the integration of endogenous, oscillatory information.

Neuromodulation has widespread potential. Investigators may elucidate functional roles of oscillatory, neural information. The role of phase and frequency information has already been documented in neuropathologies such as schizophrenia (Perez et al., 2017), epilepsy (Chu et al., 2017; Nariai et al., 2017), and Parkinson’s disease (Cozac et al., 2016; Latreille et al., 2016). Given oscillatory differences between medically induced coma, sleep and awake states, could tACS augment perioperative EEG for anesthesia dosing, confirming unconsciousness, and facilitating recovery from anesthesia? Industry may anticipate applications in online modulation of cognitive states for human-operators, as well as brain-computer interfaces. Attention-modulation would have wide-spread industrial applications by facilitating participants’ integration of environmental cues and information.

Here, we have introduced rigorously controlled tACS protocols, methodological precautions and guidelines for future tACS implementation. Recent developments in tACS scholarship, only partially presented here, foreshadow the intersection of numerous neurocognitive specializations, through the multibeneficial lens of oscillatory activity.

## Author Contributions

A.V.T. and K.Y. compiled literature for this review. K.Y. provided mentorship on methods, theory and prose. A.V.T. wrote the manuscript and provided illustrations.

